# PERK Deficiency Amplifies Molecular, Structural, and Network Vulnerability to Repetitive Mild Traumatic Brain Injury

**DOI:** 10.64898/2026.03.04.709563

**Authors:** Marangelie Criado-Marrero, Sakthivel Ravi, Daylin Barroso, Tristyn N. Garza, Diana M. Cuestas Torres, Christian Lessard, Juan Pablo Castano, MacKenzie L. Bolen, Uriel Rubinovich, Stefan Prokop, Paramita Chakrabarty, Laura P. W. Ranum, Marcelo Febo, Jose Francisco Abisambra

## Abstract

Repetitive mild traumatic brain injury (rmTBI) produces cumulative cellular stress that can lead to progressive brain dysfunction, yet the mechanisms governing vulnerability to repeated injury remain unclear. Protein kinase RNA-like endoplasmic reticulum kinase (PERK) regulates cellular proteostasis through the unfolded protein response and is implicated in neurodegeneration and acute brain injury. Here, we directly tested the role of PERK deficiency in shaping the brain’s response to rmTBI. Using a mouse model of neuronal PERK deficiency, we combined spatial proteomics and tissue analyses with resting-state functional MRI and diffusion tensor imaging to assess molecular, functional, and structural outcomes after rmTBI. PERK deficiency increased susceptibility to rmTBI-induced disruption of protein homeostasis, altered large-scale functional connectivity, and exacerbated white matter microstructural changes consistent with axonal and myelin damage. Molecular alterations were spatially aligned with imaging-defined network and white matter abnormalities. These findings identify PERK signaling as a key determinant of brain resilience to repetitive mild injury and link ER stress dysregulation to network-level dysfunction following rmTBI.

## Introduction

Traumatic brain injury (TBI) initiates both immediate mechanical disruption and delayed secondary biochemical cascades that compromise neuronal survival. These secondary processes include oxidative stress, neuroinflammation, mitochondrial dysfunction, and activation of the endoplasmic reticulum (ER) stress response, a central regulator of cellular proteostasis (reviewed in^1–3^). The unfolded protein response (UPR) is engaged to restore ER homeostasis by transiently reducing global protein synthesis while enhancing chaperone activity and protein degradation pathways (reviewed in^4^). This response is coordinated by three ER-resident stress sensors—protein kinase RNA-like endoplasmic reticulum kinase (PERK), inositol-requiring enzyme 1 (IRE1), and activating transcription factor 6 (ATF6)—which collectively determine cellular fate under proteostatic stress^5^. However, when ER stress is prolonged or recurrent, sustained UPR activation shifts from an adaptive to a pro-degenerative cascade^6^.

Among the UPR branches, chronic PERK signaling is particularly detrimental to neuronal function. Persistent PERK–eIF2α activation enforces translational repression, leading to synaptic dysfunction, impaired plasticity, and neuronal loss^7–10^. Dysregulated PERK activity has been extensively implicated in neurodegenerative disorders, including Alzheimer’s disease, multiple sclerosis, progressive supranuclear palsy (PSP), Parkinson’s disease, and tauopathies^11–16^, positioning PERK as a convergent stress node linking proteostasis failure to synaptic and network-level degeneration. However, PERK signaling is not uniformly deleterious; its impact is highly dependent on developmental stage, cell type, and duration of activation.

Transient or moderate PERK activation can support proteostatic adaptation and neuronal survival^17,18^, whereas insufficient PERK signaling, such as that conferred by hypomorphic EIF2AK3 variants, fails to prevent tau accumulation under stress and increases susceptibility to tauopathies^19,20^ and exacerbates oligodendrocyte loss^21^, highlighting a narrow functional window for PERK-dependent protection.

Emerging evidence indicates that PERK–eIF2α signaling also contributes to TBI-related neuropathology^22–24^. In experimental brain injury models, PERK activation suppresses CREB signaling and synaptic proteins such as PSD95, leading to cognitive impairments, while pharmacological PERK inhibition restores synaptic integrity and memory performance^22^. PERK signaling also promotes post-traumatic neuroinflammation through the STING–IFNβ axis, with PERK blockade attenuating cytokine induction and glial activation^25^. Consistently, pharmacological or genetic suppression of PERK reduces neuronal apoptosis and ER stress markers following brain injury and in neurodegenerative models^23,26–28^.

Despite these advances, the contribution of PERK signaling to neuronal vulnerability and post-injury inflammatory following repetitive mild TBI (rmTBI) remains incompletely understood. rmTBI is characterized by repeated subthreshold insults that cumulatively induce persistent cellular stress and progressive network dysfunction^29–31^, yet the molecular mechanisms underlying this susceptibility are poorly defined. In this study, we evaluated the impact of PERK deficiency on rmTBI-induced brain dysfunction using *in vivo* multimodal approaches. We integrated spatial protein profiling and tissue analyses with resting-state functional MRI (rsfMRI) and diffusion tensor imaging (DTI) to assess protein disruption, as well as functional connectivity and white matter microstructure as sensitive markers of axonal integrity and demyelination.

Together, these data reveal that PERK-driven ER stress mechanistically links cumulative molecular injury to systems-level network dysfunction after repetitive mild TBI, positioning PERK signaling as a pivotal regulator of brain susceptibility to repetitive injury.

## Methods

### Animals, Genotyping, and Experimental Groups

We used CRE (control) and PERK-knockdown (PKD) mice of both sexes. Animals were housed under a 12-h light/dark cycle with ad libitum access to food and water. All experimental procedures were approved by the University of Florida Institutional Animal Care and Use Committee and followed NIH guidelines. Mice were assigned to four experimental groups based on genotype and injury status: CRE-Sham (n=4), CRE-rmTBI (n=5), PKD-Sham (n=6), and PKD-rmTBI (n=7). All analyses were performed blinded to group identity. Injury was induced in mice aged 12–14 months, with behavioral assessments conducted at 4 days post-injury (dpi) and imaging at 5 dpi, followed by brain tissue collection at 7 dpi (**Experimental Timeline in Fig. 1A**).

**Figure 1.**
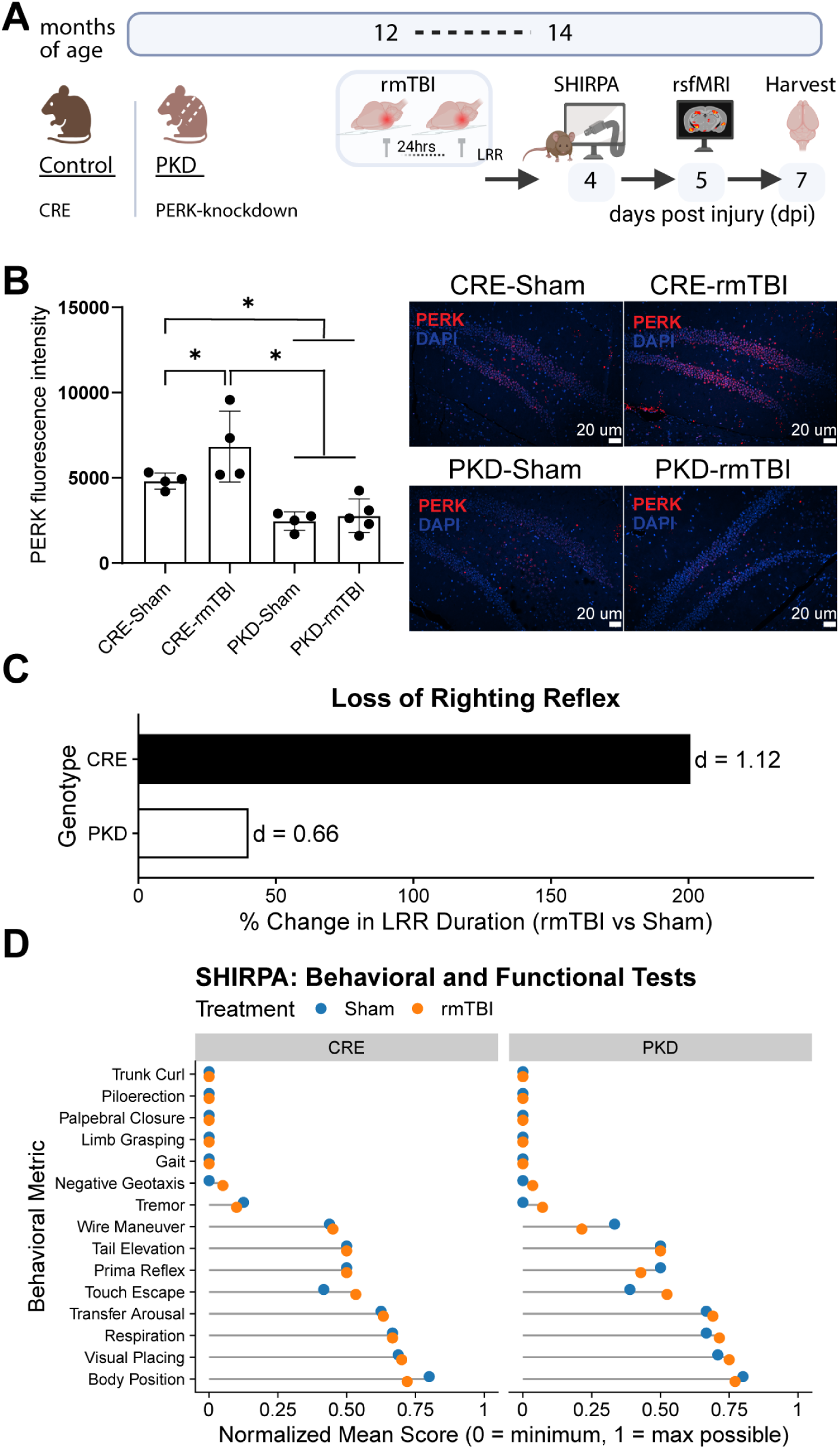
Experimental timeline and validation of PERK knockdown. **(A)** Schematic of the study timeline. CRE control (CRE) and CaMKIIα-Cre–driven PERK knockdown (PKD) mice were aged to 12–14 months and subjected to repetitive mild traumatic brain injury (rmTBI). Behavioral testing (SHIRPA) was performed at 4 days post-injury (dpi), followed by resting-state fMRI at 5 dpi and brain collection at 7 dpi. **(B)** Validation of PERK knockdown in hippocampal dentate gyrus. Representative 20x images are shown for each group. Two-way ANOVA with BH correction; *p <0.05. **(C)** Loss of righting reflex (LRR) following rmTBI. Bars represent percent change in LRR relative to Sham for each genotype. CRE-rmTBI mice showed a >200% prolongation of LRR compared with CRE-Sham (Cohen’s d = 1.12). PKD-rmTBI mice showed a moderate increase (d = 0.66). **(D)** SHIRPA behavioral assessment at 4 days post-injury. Normalized mean scores are shown for each behavioral metric in Sham (blue) and rmTBI (orange) groups. CRE-Sham (n = 4), CRE-rmTBI (n = 5), PKD-Sham (n = 6), PKD-rmTBI (n = 7).

PKD mice were generated by crossing Eif2ak3 floxed mice (PERK^loxP; Jackson Laboratory, #009340) with CaMKIIα-Cre transgenic mice (Tg(Camk2a-cre)T29-1Stl/J; Jackson Laboratory, #005359), resulting in PERK reduction in CaMKIIα-expressing excitatory neurons. Both parental lines were backcrossed onto a C57BL/6J background for >10 generations prior to experimental breeding to minimize background variability. Genotyping was performed by PCR to detect the presence of the Eif2ak3 floxed allele and the Camk2a-Cre transgene. CamKIIα-Cre–positive mice lacking the floxed PERK allele served as CRE controls (normal PERK expression), whereas CamKIIα-Cre–positive mice carrying the floxed Eif2ak3 allele were classified as PERK knockdown (PKD) animals. Functional reduction of PERK protein in PKD mice was confirmed at the protein level in brain tissue (**Fig. 1B**).

### Repetitive Mild Traumatic Brain Injury (rmTBI)

Repetitive mild TBI was induced as described in our previous studies using a closed-head impact system (Closed-Head Impact Model of Engineered Rotational Acceleration, CHIMERA) that delivers controlled dorsal cortical acceleration without skull fracture^29,30^. Mice received two mild impacts 24 h apart at 0.6 joules under brief isoflurane anesthesia (induction 3-4% and maintenance 1-2.5%). Sham animals underwent anesthesia and handling without impact. Two days of post-procedural subcutaneous meloxicam (10 mg/kg) were provided according to our IACUC guidelines.

### Neurological Assessment

Following the rmTBI, mice were placed in a supine position in the recovery chamber and the time required to recover normal posture was recorded as the loss of righting reflex (LRR). This measure indexes acute neurological suppression and is commonly used as a rodent analogue of loss-of-consciousness duration in humans^32^. Four days after injury, neurological function was assessed using the SmithKline Beecham – Harwell – Imperial College – Royal London Hospital – Phenotype Assessment (SHIRPA), which evaluates basic motor activity, reflexes, sensory responses, and simple coordination tasks^33^. Scores were derived from standardized observations across multiple domains to profile early post-injury deficits.

### MRI Acquisition

All MRI studies were performed using an 11.1T horizontal-bore scanner (Magnex Scientific, Oxford, UK) equipped with a radiofrequency coil tuned to 470.7 MHz (1 H resonance). Mice were anesthetized with isoflurane delivered in medical air (70% N₂ / 30% O₂), beginning at 4% for induction and maintained at 1% during scanning. Three imaging modalities were collected: anatomical T2-weighted MRI, resting-state functional MRI (rsfMRI), and diffusion-weighted images (DWI).

Structural MRI: T2-weighted anatomical images were acquired using a TurboRARE sequence (TE = 41.4 ms, TR = 4000 ms, RARE factor = 16, 12 averages) with a 15 × 15 mm field of view, 0.9-mm slices, and a 256 × 256 matrix, yielding an in-plane resolution of 58.6 × 58.6 μm across 14 interleaved ascending coronal slices.

Resting-state fMRI: rsfMRI data were collected using a spin-echo EPI sequence (TE = 15 ms, TR = 2000 ms, 600 volumes, matrix = 64 × 48, 14 slices) matched to the anatomical geometry. For distortion correction, two single-volume SE-EPI scans with opposite phase-encoding directions (positive and negative PE blips) were acquired using the Bruker ParaVision method provided by Dr. Matthew Budde (Medical College of Wisconsin).

Diffusion MRI: DWI data were acquired using a multi-shot SE-EPI protocol with two diffusion shells (b = 600 and 2000 s/mm²) sampled across 46 diffusion directions, plus B0 reference images. The acquisition used TE = 19.17 ms and TR = 4 s, with an in-plane resolution of 117 × 115 μm, 0.7-mm slice thickness, and a 128 × 96 matrix across 17 coronal slices. Two datasets with identical geometry were collected: a 47-volume acquisition representing the full diffusion scheme and a 7-volume acquisition used for low-b/B0 estimation.

### Image Preprocessing

All preprocessing was performed on the University of Florida HiPerGator supercomputing cluster. Bruker DICOM files were converted to NIfTI format and processed using AFNI, FSL, ANTs, and custom MATLAB^34,35^. <u>rsfMRI preprocessing</u> included distortion correction (TOPUP), motion correction (FLIRT), despiking, temporal filtering, and ICA-based denoising. Skull stripping was completed using a pulsed-coupled neural network algorithm with manual refinement in ITK-SNAP^36^. Functional scans were aligned to each mouse’s anatomical image and nonlinearly registered to a downsampled Allen Mouse CCFv3 atlas^37^. Time series were then extracted from 148 predefined regions of interest (ROIs). <u>Diffusion preprocessing</u> included denoising, Gibbs-artifact removal, susceptibility and eddy-current correction, and intensity normalization. Diffusion tensors were estimated using FSL and Camino to generate fractional anisotropy (FA), mean diffusivity (MD), radial diffusivity (RD), and axial diffusivity (AD) maps. All diffusion maps were nonlinearly registered to the Allen Mouse CCFv3 atlas, and ROIs showing misregistration, susceptibility distortion, or signal dropout were excluded. For each ROI, mean DTI values were extracted and examined to characterize microstructural properties.

### Functional Connectivity and Network Analysis

Pairwise Pearson correlations were computed between the 148 ROI time series for each subject, yielding 10,878 unique functional connections. Correlation coefficients were Fisher z-transformed before analysis. Graph-theoretic metrics were derived to assess network organization. Global efficiency quantified whole-brain integrative capacity, and network strength represented the summed connectivity across all edges. Global metrics were evaluated across 2–40% sparsity thresholds to ensure robustness, whereas regional metrics were computed at a fixed 10% threshold to balance sensitivity to strong and moderately weighted edges.

### Diffusion Tensor Imaging

Group differences in FA, MD, RD, and AD were examined across ROIs to assess microstructural alterations associated with axonal integrity and water diffusion. Statistical inference for DTI metrics followed the resampling framework described in the Statistical Analysis section.

### Spatial Proteomic Profiling

Spatial proteomic profiling was performed on 5-µm coronal Formalin-formalin fixed paraffin - embedded (FFPE) brain sections using the Bruker Spatial Biology – NanoString – Bruker GeoMx Digital Spatial Profiling platform, following manufacturer guidelines and as described in ^29^. Nuclei were labeled with SYTO13, and microglia were visualized with Iba1 morphology markers to guide ROIs selection. We sampled anatomically defined areas, including dorsal and ventral cortex, hippocampus, thalamus, corpus callosum, and the optic tract. The dorsal cortical regions (CXd) include the somatosensory and posterior parietal cortices, whereas the ventral cortical regions (CXv) include the auditory, temporal, ectorhinal, perirhinal, and piriform cortices. We quantified proteins using Neuropathology, Neurodegeneration, Glial, and Autophagy module panels (**Supplemental Table 1**). Segments passing GeoMx quality control were normalized to housekeeping proteins, corrected for background signal, and exported from GeoMx DSP analysis suite for downstream statistical analysis in R (permutation testing, effect-size estimation, and volcano plot generation).

### Immunohistochemical analysis

Microglial (Iba1) and astrocytic (GFAP) responses were assessed by chromogenic immunohistochemistry, while PERK expression was evaluated using immunofluorescence on formalin-fixed, paraffin-embedded brain tissue. Coronal sections (5 µm) were heated at 60 °C for 15 min, deparaffinized in xylene, and rehydrated through graded ethanol followed by deionized water.

For chromogenic immunohistochemistry, antigen retrieval was performed in 10 mM citrate (pH 6.0) containing 0.5% Tween-20 using a steamer for 30 min. Endogenous peroxidase activity was quenched by incubation in PBS containing 0.3% hydrogen peroxide and 10% Triton X-100. Sections were blocked with 10% goat serum at room temperature and incubated overnight at 4 °C with primary antibodies against Iba1 (1:1000; PA5-27436, Invitrogen) or GFAP (1:1000; clone GA5, #CS3670S, Cell Signaling Technology). The following day, sections were incubated with biotinylated secondary antibodies, processed using an avidin–biotin complex detection system, and developed with DAB chromogen. Immunoreactivity was quantified using ImageScope software (v12.4.3.5008) with the positive pixel count algorithm. Mean values from three sections per animal were used for statistical analyses.

For immunofluorescence, antigen retrieval was performed in 10 mM sodium citrate buffer (pH 6.0) containing 0.05% Tween-20 using a pressure cooker for 3 min. Sections were blocked for 1 h in 5% bovine serum albumin (BSA) with 0.3% Triton X-100 and incubated overnight at 4 °C with primary antibodies against PERK (rabbit polyclonal, 1:250; GTX129275, GeneTex) and FOX3/NeuN (chicken polyclonal, 1:3000; CPCA-FOX3, EnCor Biotechnology) diluted in 2.5% BSA with 0.3% Triton X-100. On the following day, sections were incubated sequentially with donkey anti-rabbit Alexa Fluor 568 and goat anti-chicken Alexa Fluor 488 secondary antibodies (1:2000 each), with PBS washes between steps. Tissue autofluorescence was quenched using ReadyProbes Tissue Autofluorescence Quenching Solution (R37630), and sections were mounted with ProLong Gold Antifade Mountant containing DAPI.

### Confocal Microscopy and Image Analysis

Fluorescent images were acquired using a Nikon Eclipse Ti2 microscope equipped with a CSU-W1 SoRa spinning-disk confocal unit. Imaging was performed using a 10× objective with excitation at 405, 488, and 561 nm. Images were captured using NIS-Elements software (version 5.42.06) and analyzed as 16-bit images in ImageJ/Fiji. Relative fluorescence intensity was calculated by subtracting mean background fluorescence (F₀), measured from regions lacking specific signal, from the measured fluorescence intensity (F − F₀). PERK signal within the dentate gyrus was quantified by an independent experimenter blinded to the experimental group.

### Statistical Analysis

All statistical analyses were performed in GraphPad (v10) and R (v4.4.2). Data distributions were inspected, and extreme outliers were removed using a predefined interquartile-range criterion under blinded group conditions. PERK immunofluorescence and LRR behavioral measures were analyzed using two-way ANOVA with genotype and injury as fixed factors. LRR magnitude is additionally reported as percentage change and Cohen’s d effect sizes. SHIRPA scores were normalized to each metric’s maximum possible value prior to statistical testing to allow comparison across behavioral domains. For spatial proteomics, log₂-transformed expression values were analyzed in limma using empirical Bayes–moderated t-tests with predefined contrasts. Volcano plots and differential expression summaries are based on BH FDR-adjusted p-values. To evaluate cross-region (local vs. widespread) protein effects, Cohen’s d effect sizes and permutation-derived p-values (15,000 iterations) were computed for each protein within each region. Proteins were classified as local (significant in a single region) or widespread (directionally consistent across multiple regions), and regional permutation p-values were combined using Fisher’s and Stouffer’s methods to assess global consistency across regions. ROI-network level and immunostaining protein-level analyses relied on permutation-derived empirical p-values. For global imaging metrics (global efficiency and network strength) and DTI measures, statistical inference was based on resampling-derived 95% confidence intervals (15,000 iterations), with significance defined by intervals excluding zero. DTI visualizations display CI ranges and directional shifts by ROI, metric, and comparison.

## Results

### PERK deficiency attenuates acute loss of righting reflex after rmTBI

To determine whether PERK reduction alters the acute response to rmTBI, we assessed neurobehavioral outcomes in middle-aged (12–14-month-old) CRE control (CRE) and CaMKIIα-Cre PERK knockdown (PKD) mice (**Fig. 1**). Efficient PERK reduction was confirmed by immunofluorescence in the dentate gyrus of the hippocampus, where PKD mice exhibited a significant decrease in PERK signal relative to CRE controls (genotype effect; F(1,14) = 29.07, p <0.0001) (**Fig. 1B**). Loss of righting reflex (LRR), a rodent correlate of transient loss of consciousness after head injury^32^, increased following rmTBI in both genotypes (injury effect; F(1,18) = 4.68, p = 0.044) (**Fig. 1C**). However, CRE mice showed a larger increase in LRR duration (200%; Cohen’s d = 1.12) than PKD mice (39%; Cohen’s d = 0.66), suggesting reduced sensitivity to acute injury in PERK-deficient animals. Broader neurological function was evaluated at 4 days post-injury using the SHIRPA phenotypic screening protocol, which provides a semi-quantitative assessment of sensorimotor coordination, reflex integrity, and arousal (**Fig. 1D**). CRE and PKD mice performed similarly across most measures. rmTBI increased touch-escape responses in both genotypes, and tremor was slightly elevated in PKD-rmTBI mice. PKD-rmTBI mice also showed a modest reduction in wire-maneuver performance and prima reflex compared to their respective controls. Together, these findings indicate that rmTBI induces mild but detectable neurobehavioral alterations, with limited genotype-specific differences.

### PERK deficiency amplifies rmTBI-induced multi-regional proteomic disruption and corpus callosum myelin vulnerability

To determine how PERK deficiency shapes region-specific protein responses to rmTBI, we performed NanoString GeoMx spatial protein profiling across six brain regions: corpus callosum (CC), dorsal cortical regions (CXd) and ventral cortical regions (CXv), hippocampus (HPC), thalamus (THL), and optic tract (OT) (**Fig. 2A**). rmTBI produced a broader and more pronounced molecular impact in PERK-deficient mice than in CRE controls (**Fig. 2B**). Across proteins altered by injury, many exhibited large effect sizes (Cohen’s d > 0.8), indicating substantial biological shifts even when statistical significance was not uniform across regions. In CRE control animals, only a limited set of proteins with large effect sizes reached statistical significance. By contrast, PKD mice exhibited multi-regional alterations in several injury-responsive proteins, including a subset affected in two or more regions. These proteins included myelin- and axonal-associated markers (*e.g.* myelin basic protein, neurofilament light), the intermediate filament Vimentin (VIM), and glial or innate immune-related proteins such as GPNMB and ApoA-I, reflecting widespread structural and inflammatory involvement under PERK deficiency. Additional multi-regional injury effects in PKD animals included increased tyrosine hydroxylase (TH) and Park7, along with decreased LRRK2, suggesting altered dopaminergic stress responses and redox/mitochondrial signaling after rmTBI. Together, these findings demonstrate that rmTBI elicits a more extensive and coordinated proteomic disturbance in PKD mice compared to CRE controls, particularly in pathways related to myelin integrity, axonal structure, dopaminergic stress responses, and glial activation.

**Figure 2.**
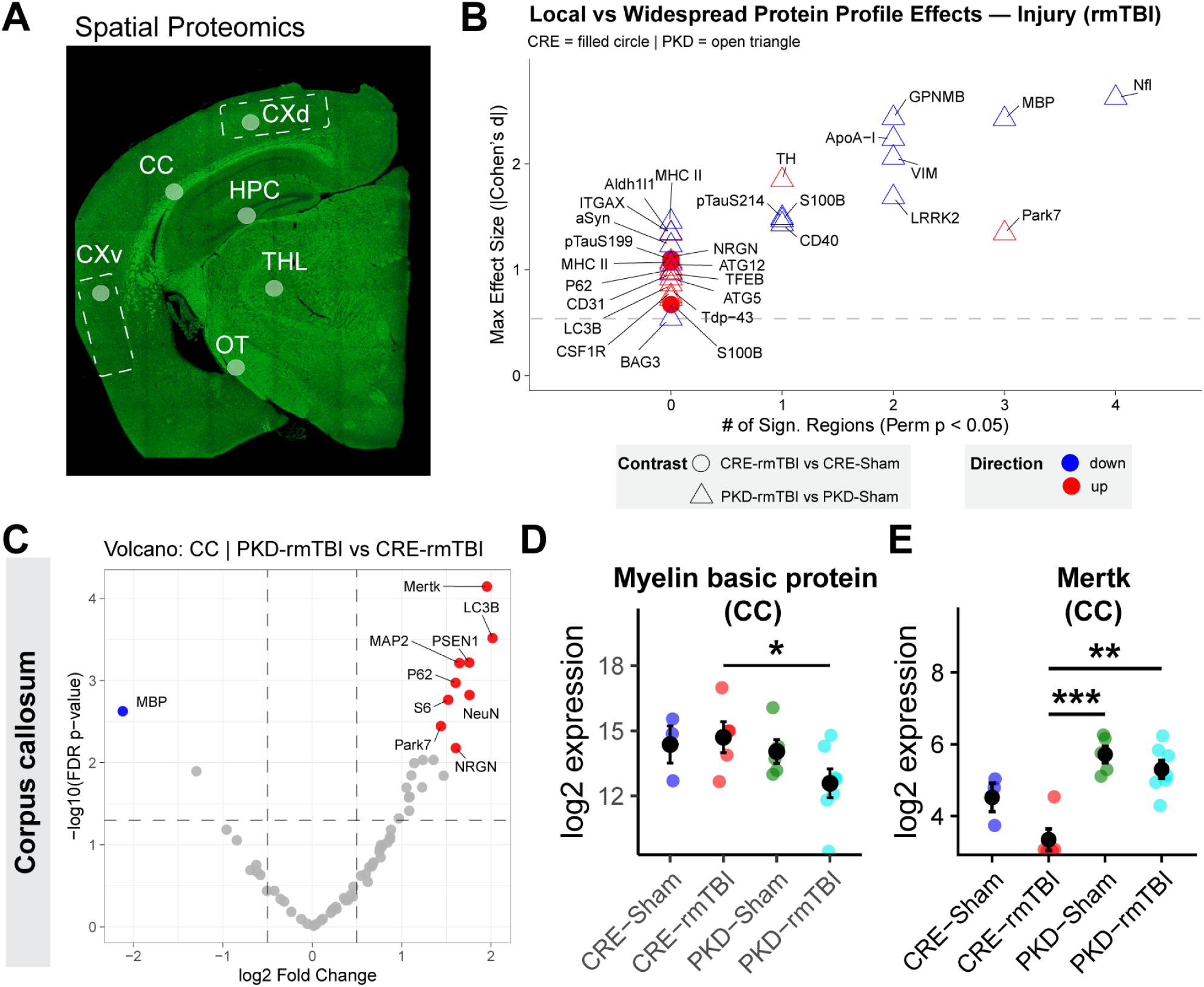
Region-specific proteomic signatures reveal myelin and axonal vulnerability after rmTBI and modulation by PERK knockdown. **(A)** Representative coronal brain section showing the six regions sampled for spatial proteomic analysis: corpus callosum (CC), Dorsal cortical regions (CXd), ventral cortex (CXv), hippocampus (HPC), thalamus (THL), and optic tract (OT). CXd and CXv boundaries varied slightly between mice based on anatomical landmarks. CXd primarily included somatosensory and posterior parietal cortices, whereas CXv comprised pooled auditory, temporal, ectorhinal, perirhinal, and piriform cortices. **(B)** Local versus widespread proteomic responses to rmTBI. Each point represents a protein, plotted by its maximum effect size (Cohen’s d) and the number of regions showing significant permutation-based (p < 0.05) changes. Filled circles denote CRE-rmTBI vs. CRE-Sham; open triangles denote PKD-rmTBI vs. PKD-Sham. Red and blue points denote increased and decreased protein levels, respectively. Myelin- and axonal-associated proteins, including Myelin Basic protein (MBP) and neurofilament light (NfL), showed the strongest regional shifts, indicating broad white-matter and axonal vulnerability after rmTBI. **(C)** Volcano plot for the corpus callosum comparing PKD-rmTBI vs. CRE-rmTBI. Changes in MBP and other injury-related targets (e.g., P62, PSEN1, LC3B, NeuN, Park7) highlight coordinated alterations in myelin integrity, neuronal markers, and inflammatory pathways under PERK deficiency. Points represent log2 fold change versus −log10(p-value); proteins with FDR < 0.05 are highlighted in red (upregulated) or blue (downregulated). **(D)** rmTBI produced a robust decrease in MBP in the corpus callosum in PKD-injured mice, consistent with white-matter sensitivity to injury. **(E)** rmTBI increased Mertk levels in corpus callosum, with an amplified response in PKD mice, suggesting heightened activation of phagocytic or debris-clearance pathways. Overall, PKD mice displayed a stronger protein shift than CRE controls. Statistical significance indicated by *p<0.05, **p<0.01, ***p<0.0001

The corpus callosum, a white-matter tract commonly disrupted after head injury^38^, showed clear rmTBI-related protein profile changes. Volcano plot analysis revealed lower myelin basic protein in PKD-rmTBI relative to CRE-rmTBI, alongside higher levels of several injury-associated markers, including LC3B, PSEN1, P62, NeuN, and Park7 (**Fig. 2C**). Group-level comparisons confirmed a significant reduction in MBP after rmTBI (**Fig. 2D**). MBP is a major structural component of myelin widely used as an indicator of white-matter integrity. Within PKD mice, rmTBI was also associated with reduced MBP levels in the HPC, OT, THL, indicating broader vulnerability of myelinated axons across affected regions. We also found that Mertk, a microglial phagocytic receptor involved in myelin and apoptotic debris clearance³¹, was increased in the corpus callosum of PKD animals, consistent with enhanced glial engagement in white-matter injury responses (**Fig. 2E**).

### Dominant genotype influence on regional proteomic profiles

Genotype exerted a stronger and more widespread influence on brain proteomic profiles than injury alone. Across regions, PERK deficiency predominantly increased protein expression, with markedly more proteins upregulated than downregulated in PKD mice under both Sham (33 up vs 2 down) and rmTBI conditions (28 up vs 7 down) (**Supplemental Fig. 1A**). Under baseline conditions, PKD mice showed elevated expression of multiple neuronal and glial proteins, including phospho-tau species, NeuN, Olig2, PSEN1, MHCII, LC3B, CD40, ITGAX, MSR1, Tau, VIM, and Mertk (**Supplemental Fig. 1B**). Many of these proteins exhibited large regional effect sizes (|Cohen’s d| ≥ 0.8) and reached significance across multiple brain regions, indicating that genotype-driven remodeling was not restricted to isolated structures but distributed across the network. Following rmTBI, genotype-dependent differences became even more regionally extensive (**Supplemental Fig. 1C**). PKD mice displayed higher levels of injury- and stress-associated proteins, including MAP2, P62, IBA1, Park7, PLA2G6, TDP-43, and Mertk, consistent with amplified cytoskeletal disruption, microglial activation, and proteostatic stress under injury. A subset of proteins, including Mertk, S6, P62, and Iba1, were elevated in PKD mice both with and without injury, suggesting that PERK deficiency establishes a primed, hyper-responsive cellular state that persists and intensifies after rmTBI.

The hippocampus showed the greatest number of genotype-dependent proteomic changes, making it the most affected region in PKD versus CRE mice (**Fig. 3**). PERK deficiency increased neuronal markers (NeuN, NRGN) and increased total Tau and phospho-Tau species (S214, S396, S404) (**Fig. 3B–G**), suggesting enhanced susceptibility of cytoskeletal and microtubule-associated pathways to rmTBI damage. PKD-Sham mice also exhibited higher APP, PSEN1, and α-synuclein levels (**Fig. 3H–J**), proteins linked to synaptic function and proteostasis, suggesting broader effects on pathways regulating protein turnover and aggregation. Calbindin, a calcium-buffering protein important for neuronal excitability, was similarly altered (**Fig. 3K**). Injured PERK mice attenuated this effect. Together, these hippocampal changes reflect the widespread genotype effects observed across regions and demonstrate that PERK knockdown shifts neuronal, axonal, and glial-associated proteomic signatures independent of rmTBI.

**Figure 3.**
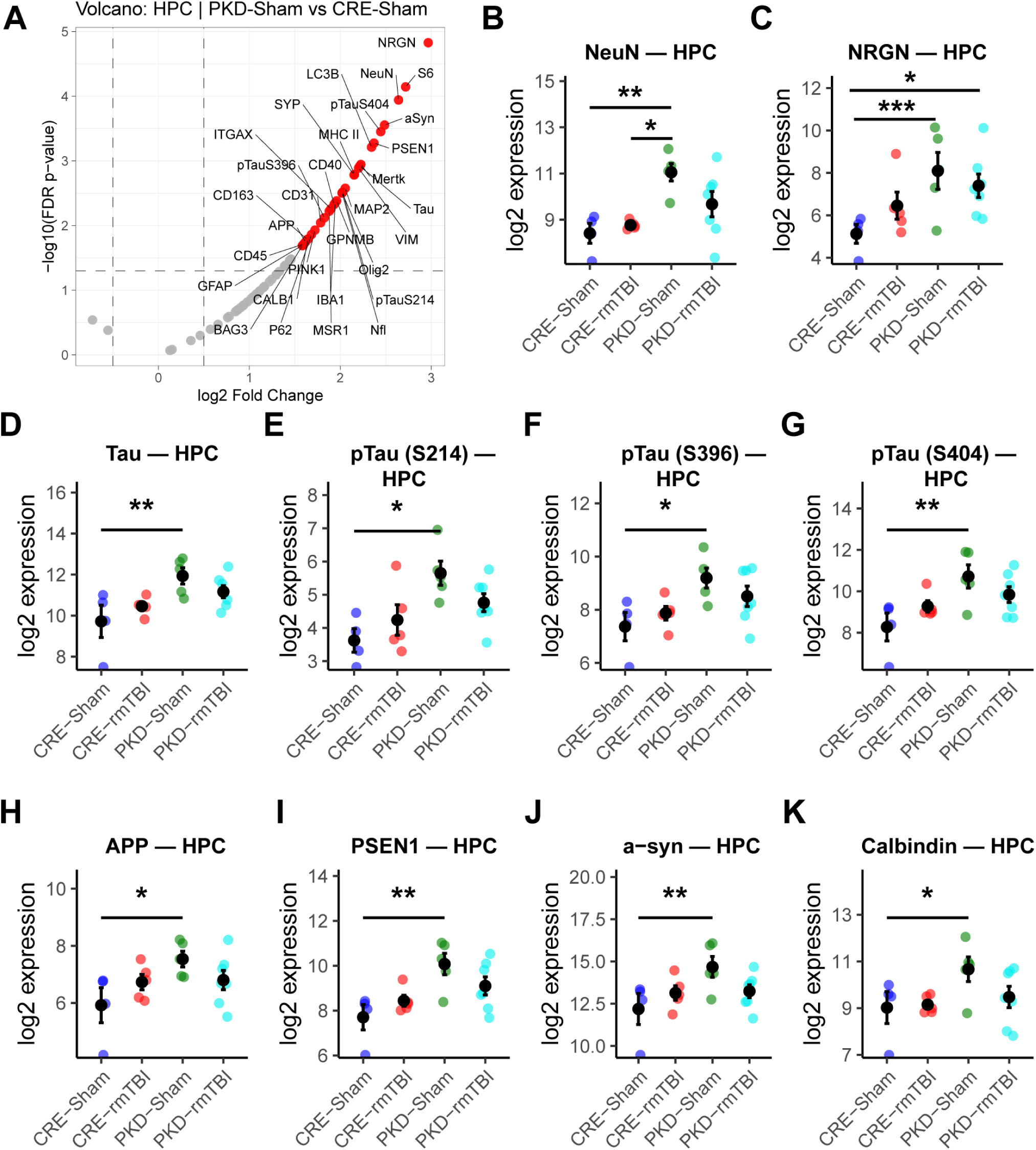
PERK deficiency produces widespread hippocampal proteomic remodeling. **(A)** Volcano plot of hippocampal proteins comparing PKD-Sham vs. CRE-Sham. Differential expression was assessed using limma with Benjamini–Hochberg FDR correction (FDR < 0.05). Red and blue points denote increased and decreased proteins, respectively. PERK deficiency increased numerous neuronal, cytoskeletal, and glial markers at baseline, including NeuN, NRGN, total Tau, phospho-Tau species, APP, PSEN1, and α-synuclein. **(B–K)** Group-wise expression of representative proteins altered by genotype. PKD mice show elevated neuronal markers **(B,** NeuN; **C**, NRGN), total Tau **(D)**, and multiple phospho-Tau species **(E–G**, S214, S396, S404). Proteostasis-related proteins **(H**, APP; **I**, PSEN1) and α-synuclein (**J**) are similarly increased in PKD animals compared to CRE. **(K)** Calbindin expression is also elevated in PKD mice. Data illustrate that PERK deficiency drives broad hippocampal molecular remodeling independent of rmTBI. HPC, hippocampus; Significance levels: *p<0.05, **p<0.01, ***p<0.0001

### Genotype and injury shape glial and autophagy responses across the hippocampus and optic tract

To contextualize the genotype-driven proteomic changes identified in the volcano analyses, we next examined glial and autophagy-related protein signatures in the hippocampus (**Fig. 4A–B**). PERK deficiency presented higher baseline levels of glial and autophagic markers relative to CRE-Sham mice. Several proteins, including CD9, CSF1R, GPNMB, ITGAX, MSR1, BAG3, BECN1, P62, PLA2G6, ULK1, and VPS35, showed more than a 2-fold increase in PKD-Sham compared to CRE-Sham. Mertk, VIM, and LC3B were among the most strongly upregulated, exhibiting approximately 4-fold higher expression in PKD animals. Following injury, PKD mice demonstrated larger reductions in key glial markers, including GPNMB, ITGAX, and VIM, than CRE mice, indicating a stronger perturbation of the glial response. In contrast, CRE-injured mice exhibited only mild upregulation of these proteins compared to CRE-Sham, consistent with a more transient or moderate inflammatory reaction.

**Figure 4.**
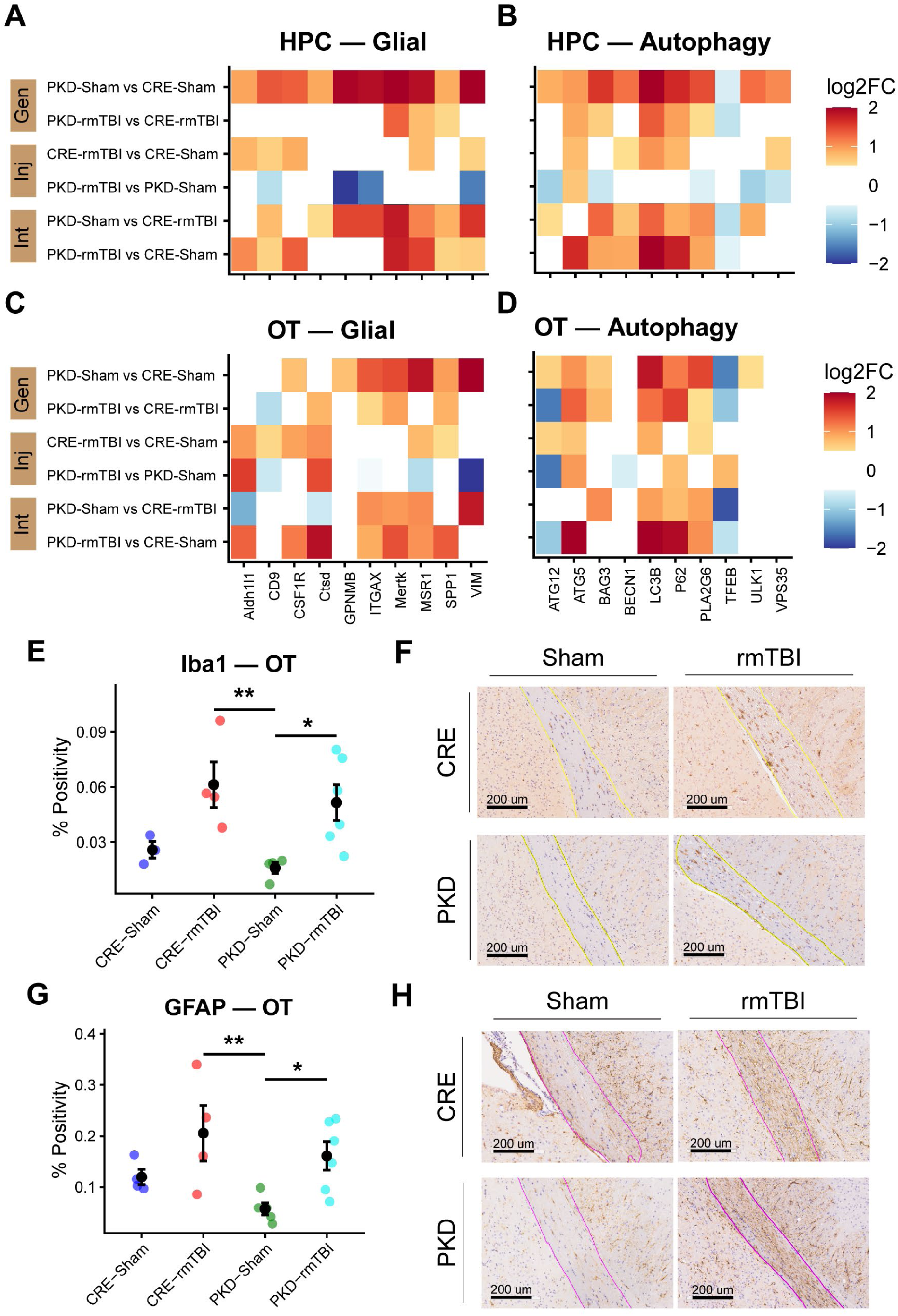
PERK deficiency enhances glial and autophagy pathway activation in hippocampus and optic tract. (A–D) Heatmaps of log2 fold changes for glial and autophagy-related proteins in the hippocampus (HPC) and optic tract (OT) across all group contrasts (Gen, genotype; Inj, injury; Int, interaction effect). PERK deficiency amplifies basal and injury-evoked glial signatures (e.g., Mertk, GPNMB, MSR1, Vimentin) and upregulates autophagy regulators (e.g., LC3B, ATG5, BAG3, P62). Quantification **(E**, Iba1; **G**, GFAP) and representative images **(F**, Iba1; **H**, GFAP; scale bars 200 µm) from optic tract immunostaining. rmTBI increased microglial and astrocytic reactivity in both genotypes. Statistical significance indicated by *p<0.05, **p<0.01

Given its long, myelinated axons being particularly susceptible to diffuse mechanical forces, the optic tract (OT) is a white matter pathway highly prone to injury. Therefore, we next assessed these signatures in the OT (**Fig. 4C-D**). Like the hippocampus, PKD-Sham mice displayed greater upregulation of glial and autophagy-related proteins relative to CRE-Sham, with VIM representing the most prominently increased protein. In both the OT and hippocampus, however, VIM levels decreased more than 50% after injury in PKD animals, suggesting a pronounced injury-induced cytoskeletal remodeling response.

Autophagy markers also showed genotype-specific patterns. TFEB, a central regulator of the autophagy–lysosomal network^39^, was reduced in PKD mice after injury. Likewise, ATG12 decreased after rmTBI in PKD animals (−0.94 log₂FC, 48% reduction in hippocampus; −1.60 log₂FC, 67% reduction in OT), whereas ATG5 increased in nearly all comparisons, indicating a divergent autophagy response in which upstream conjugation steps are enhanced while regulatory components are suppressed.

Finally, we quantified glial activation in the optic tract using Iba1 (microglia) and GFAP (astrocytes). Iba1 immunoreactivity increased by 120% in CRE mice (Cohen’s d = 1.94, p = 0.06) and by 200% in PKD mice (Cohen’s d = 2.07, p = 0.023) following rmTBI (**Fig. 4E–F**). Astrocytic responses followed a similar pattern: CRE-rmTBI mice exhibited a 75% increase in GFAP signal (Cohen’s d = 1.08, p = 0.13), whereas PKD-rmTBI mice showed a 160% increase (Cohen’s d = 2.0, p = 0.031) relative to their respective sham controls (**Fig. 4G–H**). These data aligned with our previous findings showing that repeated mild TBIs elicit a robust neuroinflammatory response in this vulnerable white matter tract^29,30^.

### PERK deficiency lowers baseline network integration and prevents the injury-associated increase in global efficiency

Given the broad proteomic divergence between genotypes, we next examined whether PERK deficiency altered network-level functional connectivity. Mean connectivity maps revealed a marked reduction in whole-brain connectivity strength in PKD mice compared with CRE controls under both Sham and rmTBI conditions (**Fig. 5A**). PKD networks showed fewer highly connected nodes and a shift toward weaker, more spatially diffuse connectivity patterns.

**Figure 5.**
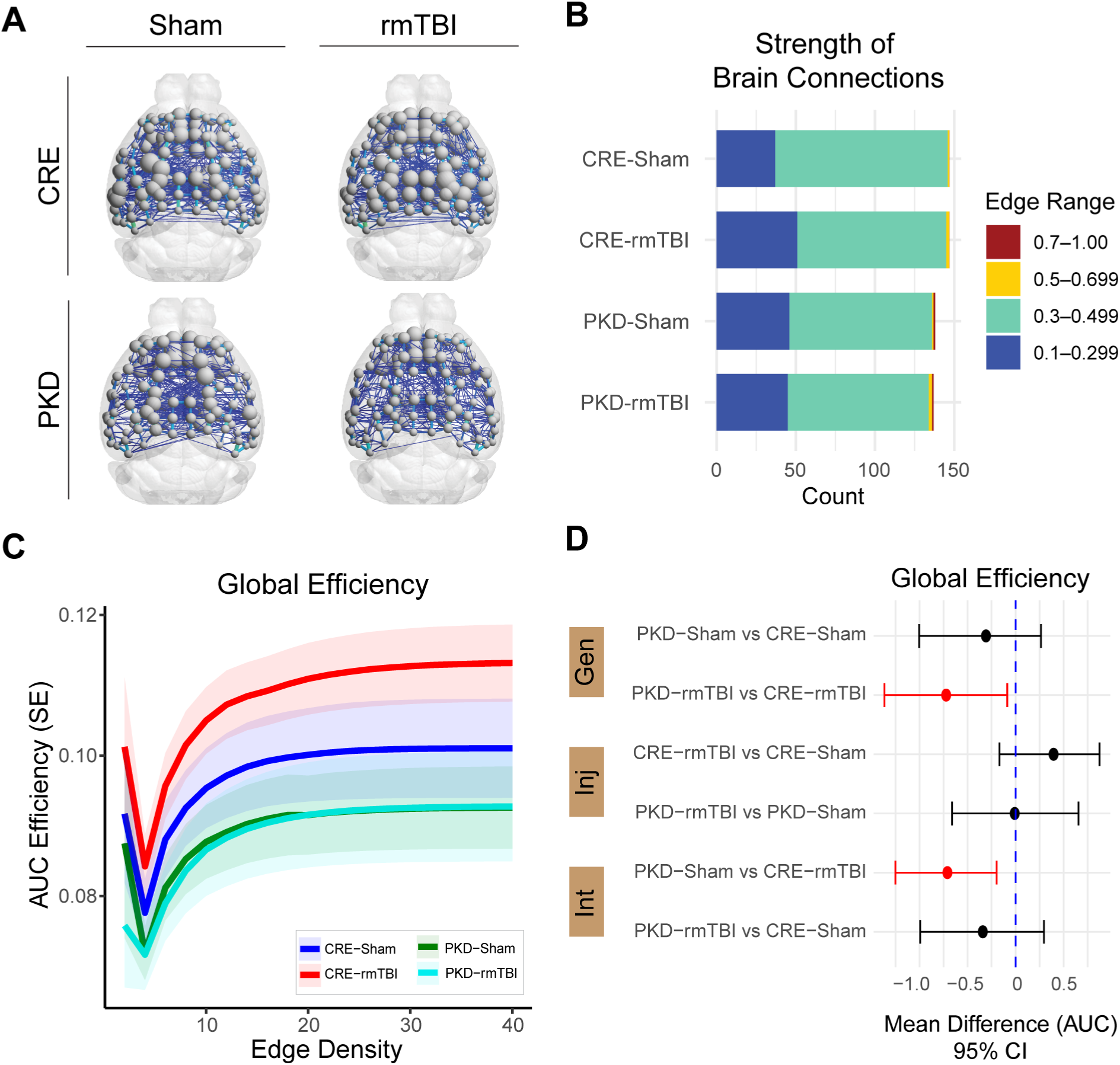
PERK deficiency weakens whole-brain connectivity and reduces global network efficiency. **(A)** Resting-state functional connectivity maps showing node strength (sphere size proportional to total connectivity per region) for CRE and PKD mice under Sham and rmTBI conditions. PKD brains exhibit fewer highly connected nodes and an overall reduction in connectivity strength. **(B)** Distribution of functional connection strengths (edge weights). PKD mice show a higher proportion of weak edges (Pearson r = 0.1–0.3) and fewer strong edges (Pearson r ≥ 0.5) relative to CRE animals, indicating reduced network robustness. **(C)** Global network efficiency curves across edge density thresholds (2–40% of strongest functional network connections). CRE-injured mice display consistently higher efficiency than PKD mice. Shaded areas represent ±SEM. **(D)** Area-under-the-curve (AUC) comparisons for global efficiency in panel **C**, shown as mean differences with 95% confidence intervals derived from bootstrap resampling (15,000 iterations). PKD mice exhibit significantly lower efficiency than CRE controls under both Sham and rmTBI conditions. Red markers denote contrasts showing significant 95% Confidence Interval (CI).

Consistent with this, PKD mice displayed a higher proportion of weak edges (Pearson’s r = 0.1–0.3) and fewer moderate edges (Pearson’s r ≥ 0.5) relative to CRE animals (**Fig. 5B**). Here, edges represent pairwise functional correlations between brain regions. This imbalance was present at baseline and persisted after rmTBI, indicating a genotype-driven vulnerability in maintaining robust functional connections. Global efficiency analyses showed a similar pattern. Only CRE mice exhibited an injury-induced increase in efficiency, a compensatory response commonly observed after diffuse brain injury^40^ PKD mice failed to mount this response, resulting in a significant global efficiency deficit compared with CRE-rmTBI mice (**Fig. 5C-D**). Together, these findings indicate that PERK deficiency weakens large-scale functional connectivity and impairs the brain’s ability to adapt network architecture following rmTBI.

Network strength reflects how robustly each region communicates with the rest of the brain, serving as an index of functional integration and resilience. Because global efficiency depends on the strength of connections across the brain, we next examined whether PERK deficiency also alters the overall distribution of network strength. Network-strength analyses showed the same pattern observed in global connectivity: PKD mice exhibited consistently lower regional strength than CRE-injured animals (**Fig. 6A-B**), indicating weaker integration of individual connections into the broader functional network. Across 148 ROIs, permutation testing confirmed that genotype was the dominant driver of these effects, with fewer regions showing genotype–injury interactions (**Fig. 6C**). Hippocampal analyses mirrored this pattern with significantly reduced node strength and node efficiency in PKD mice, regardless of injury, indicating impaired information processing within this hub region (**Fig. 6D**). Node degree decreased only in the genotype×injury interaction, suggesting that rmTBI further disrupts the number of functional connections specifically in PKD animals. Overall, PERK deficiency produces widespread reductions in regional connectivity, with the hippocampus showing marked deficits in both the strength and efficiency of functional interactions and additional injury-specific disruptions in network topology.

**Figure 6.**
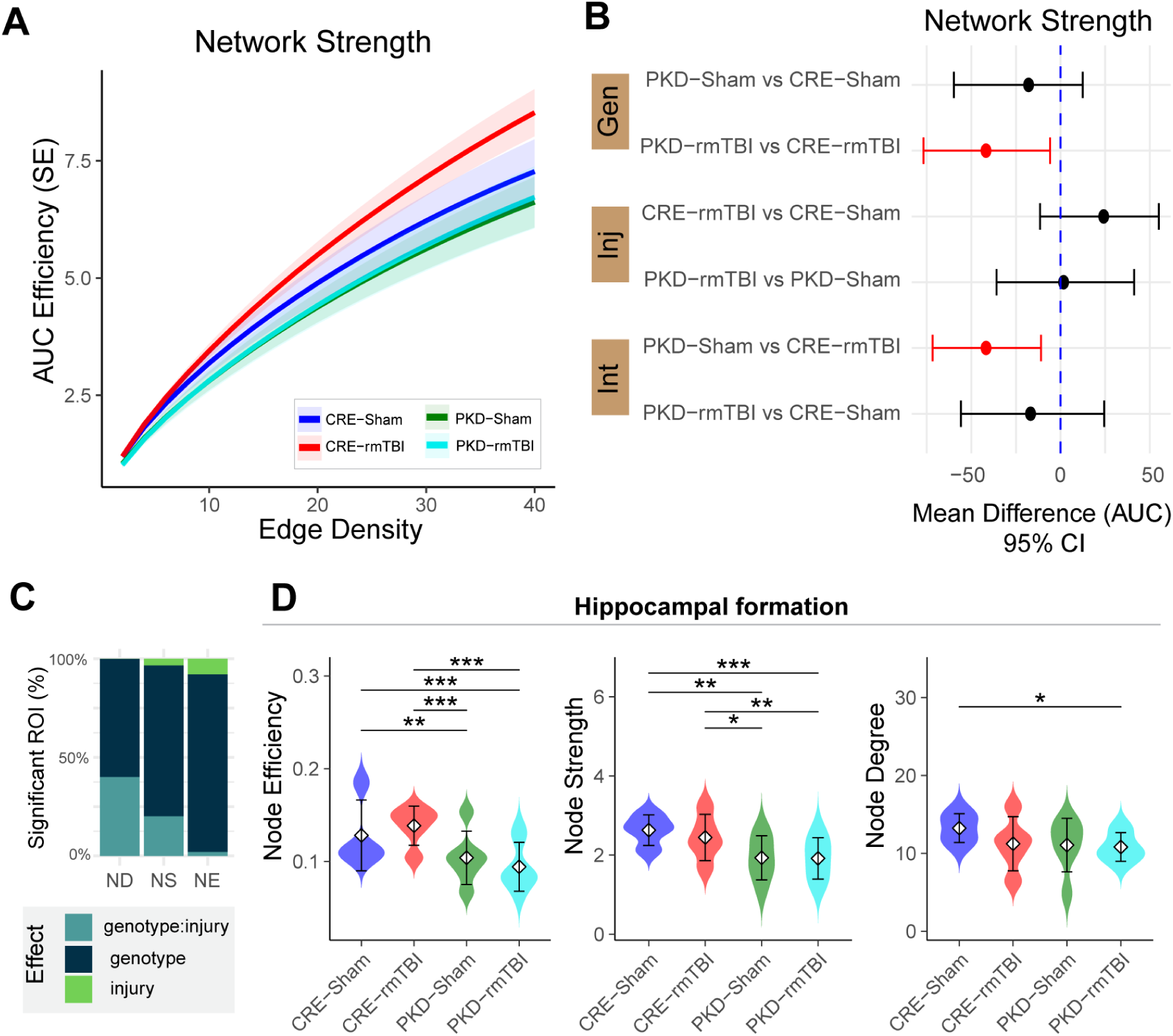
PERK deficiency reduces global network strength and alters hippocampal node-level connectivity. **(A)** Network strength curves across edge densities for each group. PKD mice show consistently lower network strength than CRE controls, with rmTBI producing an additional reduction in both genotypes. **(B)** Network-strength area-under-the-curve (AUC) differences between groups shown as mean differences with 95% confidence intervals derived from bootstrap resampling (15,000 iterations). Red markers indicate contrasts for which the 95% CI does not include zero. Confidence intervals indicate significant decreases in PKD vs. CRE mice under both Sham and rmTBI conditions (red), confirming a genotype-driven reduction in global connectivity. **(C)** Percent of whole brain regions (ROI, n = 148) showing significant effects for node degree (ND), node strength (NS), and node efficiency (NE). Genotype and genotype×injury interactions account for the majority of node-level differences. **(D)** Violin plots of hippocampal node metrics. Data presented in mean ± SD. PERK deficiency lowers node efficiency and node strength at baseline, with rmTBI further reducing connectivity in both genotypes. Node degree shows a modest but significant decrease in PKD mice. Together, these results demonstrate that PERK loss impairs both global network integration and regional node-level connectivity. Statistical significance *p<0.05, **p<0.01, ***p<0.0001

### PERK deficiency and rmTBI reduce white-matter microstructural integrity

We next investigated whether these functional network abnormalities were accompanied by microstructural changes detectable with diffusion MRI (**Fig. 7**). Fractional anisotropy (FA) was reduced by both injury and genotype. In CRE mice, rmTBI decreased FA in cortex, hippocampus, and fornix, consistent with axonal disruption. PKD mice also showed lower FA than CRE-Sham controls, indicating baseline microstructural weakness. When genotype and injury were combined (PKD-rmTBI vs. CRE-Sham), FA reductions appeared across additional regions, including the optic tract, suggesting that PERK deficiency increases the extent of white matter disruption after rmTBI. rmTBI had minimal impact on PKD mice, except for reduced mean and axial diffusivity in the optic tract. Thus, PERK deficiency and rmTBI each contribute to reduced white matter integrity.

**Figure 7.**
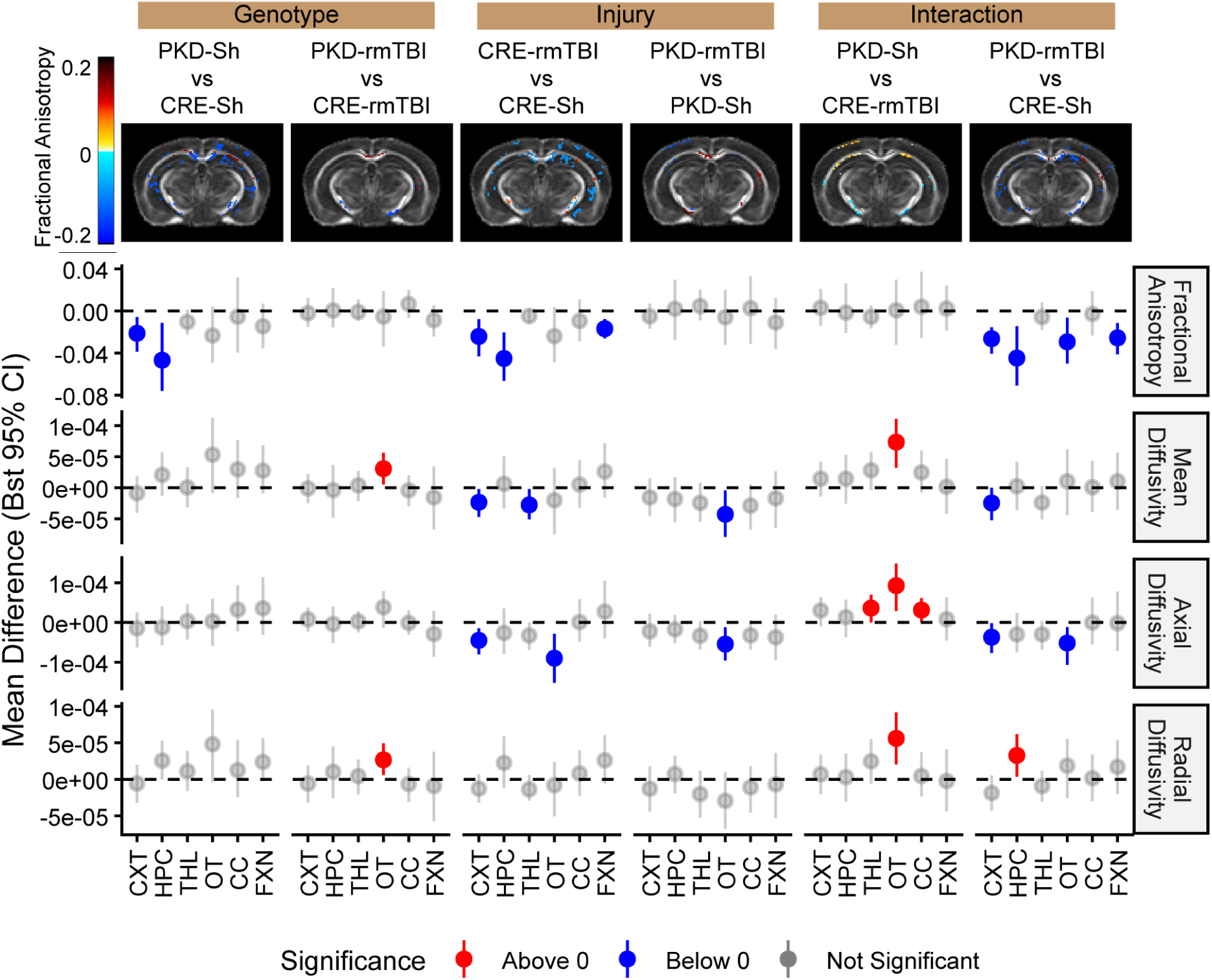
PERK deficiency and rmTBI alter white-matter microstructural integrity. Bootstrapped (Bst) mean differences (95% confidence intervals) for fractional anisotropy (FA), mean diffusivity (MD), axial diffusivity (AD), and radial diffusivity (RD) across white- and gray-matter regions. Points represent bootstrapped mean differences, and error bars denote 95% confidence intervals (15,000 iterations). Red points indicate significant positive differences (CI > 0), blue points indicate significant negative differences (CI < 0), and gray points denote non-significant comparisons (CI includes 0). The dashed horizontal line indicates no difference (0). rmTBI reduced FA in CRE mice, particularly in hippocampus and cortex, consistent with axonal disruption. PKD mice exhibited lower baseline FA relative to CRE-Sham controls. Combined genotype and injury (PKD-rmTBI vs. CRE-Sham) was associated with broader reductions in FA across additional regions, including optic tract. In contrast, rmTBI increased RD in PKD mice within the corpus callosum, suggesting region-specific alterations in myelin-sensitive metrics.

## Discussion

Our findings suggest that PERK signaling plays a critical role in shaping brain vulnerability to rmTBI by regulating how cellular stress is translated into structural and network-level dysfunction. PERK deficiency did not exacerbate overt acute behavioral impairment, as reflected by attenuated loss of righting reflex responses. Instead, it shifted the progression of injury toward greater white matter vulnerability and impaired functional network organization during the acute post-injury period. This dissociation underscores that early physiological severity does not necessarily predict later network integrity when proteostatic regulation is disrupted. Rather than uniformly worsening injury severity, PERK deficiency modifies how injury-related stress accumulates and propagates across brain systems over time.

At baseline, PERK deficiency established a genotype-dependent molecular landscape characterized by broad hippocampal proteomic remodeling. PERK-deficient mice exhibited increased neuronal and dopaminergic-related markers, including NeuN, NRGN, TH, and Park7, consistent with prior studies showing that reduced PERK signaling can support neuronal health and dopaminergic pathway integrity^13,16,41^. However, this apparent neuronal benefit coexisted with altered stress-handling capacity. Spatially resolved protein profiling showed coordinated alterations in proteins reflecting myelin integrity (MBP), axonal injury (NfL), cytoskeletal remodeling (Vimentin), innate immune activation (GPNMB, ApoA-I), and oxidative stress (PINK1, BAG3), which are established markers of white-matter injury and neurodegenerative vulnerability in TBI^42–44^. These changes likely represent secondary consequences of impaired proteostatic regulation rather than direct PERK-dependent control.

A key feature of the PERK-deficient phenotype was elevated baseline expression of glial and autophagy-related proteins, particularly in the hippocampus and optic tract, consistent with a pre-stressed state. Autophagy regulators (LC3B, ATG5, P62) and glial markers (GPNMB, Vimentin, Mertk) are commonly upregulated to maintain homeostasis under chronic stress^43^ (reviewed in^45^). However, rmTBI exposed the limits of this adaptation: following injury, these pathways were reduced rather than further induced in PKD mice, in contrast to control animals. This pattern differs from conditions such as Alzheimer’s disease, where sustained PERK activation is associated with persistent autophagic P62 accumulation^28^. Further evidence of impaired autophagy regulation was provided by divergent changes in ATG12 and ATG5 expression. Because the ATG12–ATG5 conjugation system is required for autophagosome formation^46^, reduced ATG12 alongside increased ATG5 indicates dyscoordination of autophagy flux rather than effective activation. This interpretation is reinforced by reduced TFEB and CD9 levels in PKD mice after injury in the hippocampus and optic tract. TFEB is a master regulator of lysosomal and autophagic gene expression^39^, while CD9 is associated with microglial activation and myelin repair^47^. Together, these changes suggest that PERK deficiency limits the capacity to maintain coordinated autophagy and glial remodeling programs under repeated stress, resulting in exhaustion of adaptive responses rather than enhanced inflammation.

Hippocampal protein profiling also revealed injury-dependent alterations in tau and related pathological proteins that further support increased molecular vulnerability in PERK-deficient mice. Following rmTBI, PKD animals exhibited increased total tau and multiple phosphorylated tau species (pTau S214, S396, S404), together with increased APP, PSEN1, and α-synuclein. These proteins are established markers of axonal stress, disrupted proteostasis, and neurodegenerative risk after brain injury^19,48,49^. Importantly, phosphorylation at these tau epitopes is commonly associated with cytoskeletal instability and impaired axonal transport rather than mature tau aggregation, indicating stress-related signaling rather than overt tauopathy. In the context of impaired PERK-mediated proteostatic control, these changes suggest that repetitive injury lowers the threshold for tau and amyloid-associated pathway engagement in hippocampal circuits.

These molecular vulnerabilities were mirrored by structural imaging findings. PERK-deficient mice exhibited lower fractional anisotropy at baseline, indicating weakened white-matter integrity prior to injury. Following rmTBI, FA reductions in CRE mice were confined to cortex, hippocampus, and fornix. We were expecting to see damage in the optic tract since this is a vulnerable area after injury^29,30^; however, injury-related deficits extended into the optic tract only under genotype and injury interaction. This expansion suggests that PERK deficiency lowers the threshold for white-matter damage following repetitive injury. The reduced global efficiency and network strength can be due to the white matter vulnerability, such as the corpus callosum with reduced MBP levels, together with increased expression of the phagocytic receptor Mertk. This indicates an elevated burden of myelin damage and increased demand for debris clearance.

Because Mertk mediates microglial engulfment of damaged myelin^50^, its upregulation suggests enhanced phagocytic activity in response to myelin disruption rather than effective repair. In the context of PERK deficiency, these changes are consistent with impaired proteostatic capacity and disrupted oligodendrocyte homeostasis, aligning with prior studies implicating PERK signaling in myelin maintenance and stress adaptation^51^.

At the systems level, PERK deficiency compromised global network architecture, reducing network strength and efficiency at baseline and amplifying rmTBI induced connectivity loss, with the hippocampus emerging as a central hub of vulnerability. This disruption links ER stress dysregulation to white matter fragility and impaired large scale brain integration. These experiments were performed in 14-month-old mice, a stage associated with emerging age-related white matter decline and reduced network efficiency^30^. Sustained PERK reduction therefore likely adds to existing age dependent circuit vulnerability. In contrast to studies showing that acute PERK inhibition can mitigate synaptic or inflammatory outcomes after injury^52^, our data demonstrate that sustained reduction of PERK signaling erodes baseline network resilience and lowers the threshold for cumulative injury. Together, these findings identify PERK signaling as a context dependent determinant of brain resilience that supports neuronal maintenance under steady state conditions and preserves network integrity, coordinated stress responses, and white matter repair during repetitive mild traumatic injury.

## Acknowledgements

The UF Neuromedicine Human Brain and Tissue Bank (HBTB) is supported by the Evelyn F. and William L. McKnight Brain Institute, the Center for Translational Research in Neurodegenerative Diseases and the 1Florida ADRC (P30 AG047266). S.P. is supported by the Charlotte and Howard Zimmerman rising star professorship at the Norman Fixel Institute for Neurological diseases. Timeline in Figure 1 Created in BioRender. Criado-Marrero, M. (2026) https://BioRender.com/x4im9zd

## Author contributions

M.C.-M: Conceptualization, Methodology, Investigation, Formal analysis, Visualization, Validation, Supervision, Funding acquisition, Writing – original draft, Writing – review & editing. S.R: Conceptualization, Methodology, Investigation, Data curation, Validation, Supervision, Writing – review & editing.

D.B: Methodology, Investigation, Data curation, Project administration. T.N.G: Data curation, Writing – review & editing.

D.M.C.-T: Data curation, Writing – review & editing.

C.L: Data curation, Formal analysis, Visualization, Writing – review & editing. J.P.C: Formal analysis, Visualization.

M.L.B.: Investigation, Methodology, Data curation, Writing – review & editing. U.R.: Methodology, Data curation, Formal analysis.

S.P.: Resources, Writing – review & editing. P.C.: Writing – review & editing.

L.P.W.R.: Writing – review & editing.

M.F.: Writing – review & editing, Resources, Methodology.

J.F.A: Writing – review & editing, Validation, Supervision, Resources, Project administration, Funding acquisition, Conceptualization.

## Data Availability Section

The datasets generated and analyzed during this study are available from the corresponding author upon reasonable request, in accordance with institutional and NIH data-sharing policies.

## Declaration for Conflict of Interest

J.F.A. serves as a consultant for Curie Bio.

**Declaration for AI tools used, if any: If needed.**

## Supplementary Materials

**Supplementary Fig. 1.**
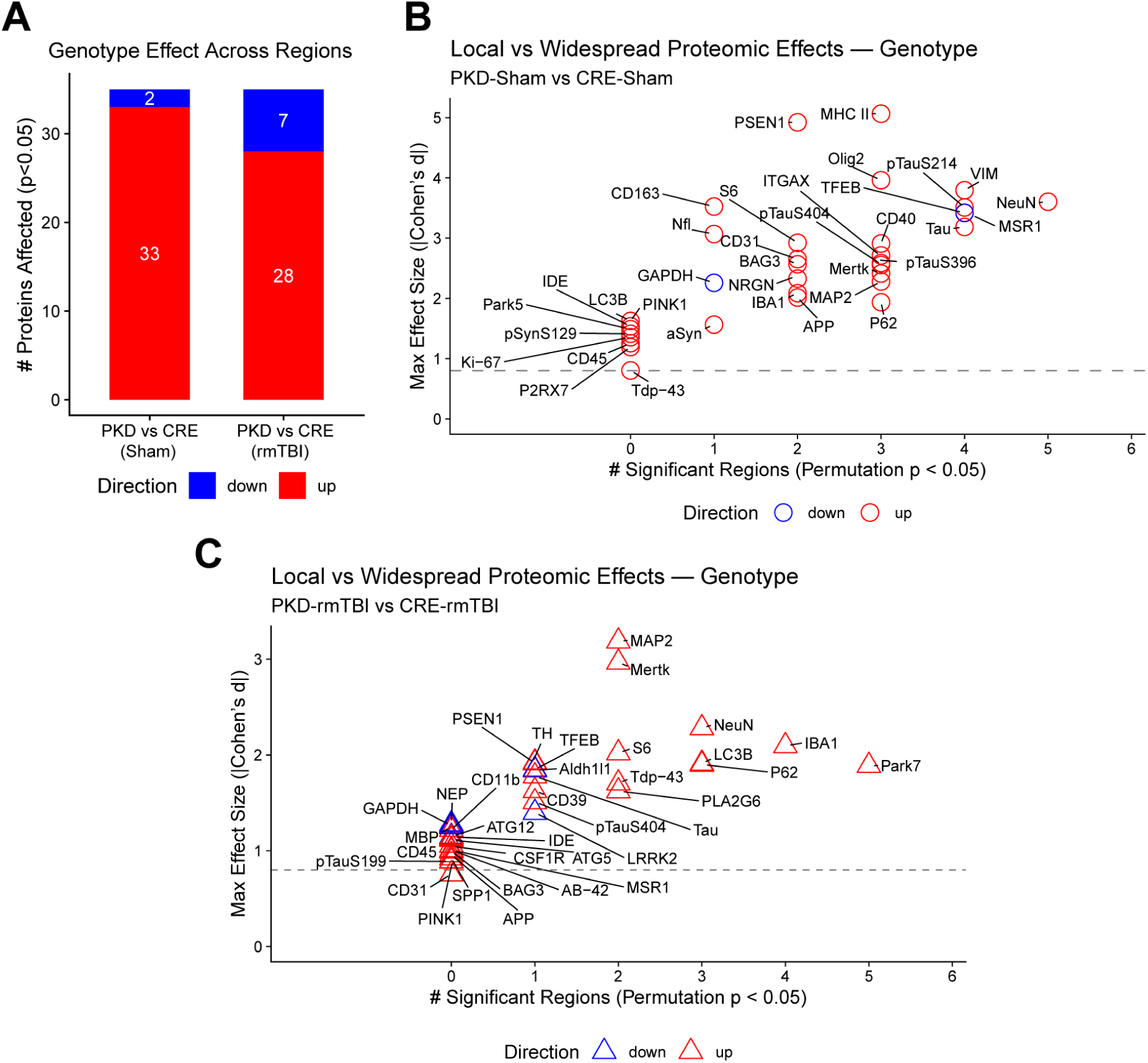
Genotype-dependent proteomic alterations across brain regions. **(A)** Number of proteins significantly altered between PKD and CRE mice under Sham and rmTBI conditions (permutation p < 0.05). Red bars indicate proteins upregulated in PKD relative to CRE; blue bars indicate downregulated proteins. **(B–C)** Local versus widespread proteomic effects across brain regions for genotype contrasts. The x-axis represents the number of brain regions in which a protein reached permutation significance (p < 0.05). The y-axis shows the maximum absolute regional effect size (|Cohen’s d|). The dashed horizontal line indicates a large effect size threshold (|d| = 0.8). Proteins appearing further to the right reflect more widespread regional alterations, whereas higher positions indicate stronger regional magnitude differences.

**Supplemental Table 1.** Proteins included in the Bruker Spatial Proteomics Panel.

|  | Bruker Spatial Panel | Protein Name | Abbrev |
| --- | --- | --- | --- |
| 1. | Alzheimer's Pathology | Apolipoprotein E | APOE |
| 2. | Alzheimer's Pathology | Amyloid Precursor Protein | APP |
| 3. | Alzheimer's Pathology | Amyloid-Beta 1-42 | AB-42 |
| 4. | Alzheimer's Pathology | P2X purinoceptor | P2RX7 |
| 5. | Alzheimer's Pathology | Phospho-Tau (S404) | pTauS404 |
| 6. | Alzheimer's Pathology | Microtubule-associated protein Tau | Tau |
| 7. | Alzheimer's Pathology | TAR DNA-binding protein 43 | Tdp-43 |
| 8. | Alzheimer's Pathology | Ubiquitin | Ub |
| 9. | Alzheimer's Pathology Extended | Beta-secretase 1 | BACE1 |
| 10. | Alzheimer's Pathology Extended | Insulin-degrading enzyme | IDE |
| 11. | Alzheimer's Pathology Extended | Neurogranin | NRGN |
| 12. | Alzheimer's Pathology Extended | Neprilysin | NEP |
| 13. | Alzheimer's Pathology Extended | Presenilin-1 | PSEN1 |
| 14. | Alzheimer's Pathology Extended | Phospho-Tau (S199) | pTauS199 |
| 15. | Alzheimer's Pathology Extended | Phospho-Tau (S214) | pTauS214 |
| 16. | Alzheimer's Pathology Extended | Phospho-Tau (S396) | pTauS396 |
| 17. | Alzheimer's Pathology Extended | Phospho-Tau (T231) | pTauT231 |
| 18. | Autophagy | Ubiquitin-like protein ATG12 | ATG12 |
| 19. | Autophagy | Autophagy protein 5 | ATG5 |
| 20. | Autophagy | BAG family molecular chaperone regulator 3 | BAG3 |
| 21. | Autophagy | Beclin-1 | BECN1 |
| 22. | Autophagy | Microtubule-associated protein 1 light chain 3b | LC3B |
| 23. | Autophagy | Sequestosome-1 | P62 |
| 24. | Autophagy | Calcium-independent phospholipase A2 | PLA2G6 |
| 25. | Autophagy | BHLH domain-containing protein | TFEB |
| 26. | Autophagy | Serine/threonine-protein kinase ULK1 | ULK1 |
| 27. | Autophagy | Vacuolar protein sorting-associated protein 35 | VPS35 |
| 28. | Glial Cell Subtyping | Cytosolic 10-formyltetrahydrofolate dehydrogenase | Aldh1l1 |
| 29. | Glial Cell Subtyping | CD9 antigen | CD9 |
| 30. | Glial Cell Subtyping | Macrophage colony-stimulating factor 1 receptor | CSF1R |
| 31. | Glial Cell Subtyping | Cathepsin D | Ctsd |
| 32. | Glial Cell Subtyping | Transmembrane glycoprotein NMB | GPNMB |
| 33. | Glial Cell Subtyping | Integrin alpha-X | ITGAX |
| 34. | Glial Cell Subtyping | Macrophage scavenger receptor types I and II | MSR1 |
| 35. | Glial Cell Subtyping | Tyrosine-protein kinase Mer | Mertk |
| 36. | Glial Cell Subtyping | SPP1 | SPP1 |
| 37. | Glial Cell Subtyping | Vimentin | VIM |
| 38. | Neural Cell Profiling | CD11b | CD11b |
| 39. | Neural Cell Profiling | Scavenger receptor cysteine-rich type 1 protein M130 | CD163 |
| 40. | Neural Cell Profiling | Platelet endothelial cell adhesion molecule | CD31 |
| 41. | Neural Cell Profiling | Ectonucleoside triphosphate diphosphohydrolase 1 | CD39 |
| 42. | Neural Cell Profiling | Tumor necrosis factor receptor superfamily member 5 | CD40 |
| 43. | Neural Cell Profiling | Protein-tyrosine-phosphatase | CD45 |
| 44. | Neural Cell Profiling | Macrosialin | CD68 |
| 45. | Neural Cell Profiling | Glyceraldehyde-3-phosphate dehydrogenase | GAPDH |
| 46. | Neural Cell Profiling | Glial fibrillary acidic protein | GFAP |
| 47. | Neural Cell Profiling | Histone H3 | H3 |
| 48. | Neural Cell Profiling | IBA1 | IBA1 |
| 49. | Neural Cell Profiling | Proliferation marker protein Ki-67 | Ki-67 |
| 50. | Neural Cell Profiling | Microtubule-associated protein | MAP2 |
| 51. | Neural Cell Profiling | H-2 class II histocompatibility antigen, Ab chain | MHC II |
| 52. | Neural Cell Profiling | Myelin basic protein | MBP |
| 53. | Neural Cell Profiling | RNA binding protein fox-1 homolog 3 | NeuN |
| 54. | Neural Cell Profiling | Neurofilament light polypeptide | Nfl |
| 55. | Neural Cell Profiling | Oligodendrocyte transcription factor 2 | Olig2 |
| 56. | Neural Cell Profiling | Rb IgG | Rb IgG |
| 57. | Neural Cell Profiling | Rt IgG2a | Rt IgG2a |
| 58. | Neural Cell Profiling | Rt IgG2b | Rt IgG2b |
| 59. | Neural Cell Profiling | Protein S100-B | S100B |
| 60. | Neural Cell Profiling | 40S ribosomal protein S6 | S6 |
| 61. | Neural Cell Profiling | Synaptophysin | SYP |
| 62. | Neural Cell Profiling | Transmembrane protein 119 | TMEM119 |
| 63. | Parkinson's Pathology | Alpha-synuclein | aSyn |
| 64. | Parkinson's Pathology | Apolipoprotein A-I | ApoA-I |
| 65. | Parkinson's Pathology | Calbindin | CALB1 |
| 66. | Parkinson's Pathology | Protein kinase domain-containing protein | LRRK2 |
| 67. | Parkinson's Pathology | Serine/threonine-protein kinase PINK1, mitochondrial | PINK1 |
| 68. | Parkinson's Pathology | Ubiquitin carboxyl-terminal hydrolase isozyme L1 | Park5 |
| 69. | Parkinson's Pathology | Parkinson disease protein 7 homolog | Park7 |
| 70. | Parkinson's Pathology | Phospho-Alpha-synuclein (S129) | pSynS129 |
| 71. | Parkinson's Pathology | Tyrosine Hydroxylase | TH |

This table lists the proteins profiled using the Bruker Spatial Biology platform. Proteins are organized by panel classification as defined by the manufacturer. Columns indicate the panel category, full protein name, and abbreviated symbol used throughout the manuscript.

